# Copy number signatures predict chromothripsis and associate with poor clinical outcomes in patients with newly diagnosed multiple myeloma

**DOI:** 10.1101/2020.11.24.395939

**Authors:** Kylee H Maclachlan, Even H Rustad, Andriy Derkach, Binbin Zheng-Lin, Venkata Yellapantula, Benjamin Diamond, Malin Hultcrantz, Bachisio Ziccheddu, Eileen Boyle, Patrick Blaney, Niccolò Bolli, Yanming Zhang, Ahmet Dogan, Alexander Lesokhin, Gareth Morgan, Ola Landgren, Francesco Maura

## Abstract

Chromothripsis is detectable in 20-30% of newly diagnosed multiple myeloma (NDMM) patients and is emerging as a new independent adverse prognostic factor. In this study, we interrogate 752 NDMM patients using whole genome sequencing (WGS) to study the relationship of copy number (CN) signatures to chromothripsis and show they are highly associated. CN signatures are highly predictive of the presence of chromothripsis (AUC=0.90) and can be used to identify its adverse prognostic impact. The ability of CN signatures to predict the presence of chromothripsis was confirmed in a validation series of WGS comprised of 235 hematological cancers (AUC=0.97) and an independent series of 34 NDMM (AUC=0.87). We show that CN signatures can also be derived from whole exome data (WES) and using 677 cases from the same series of NDMM, we were able to predict both the presence of chromothripsis (AUC=0.82) and its adverse prognostic impact. CN signatures constitute a flexible tool to identify the presence of chromothripsis and is applicable to WES and WGS data.

## Introduction

Chromothripsis, a catastrophic chromosomal shattering event associated with random rejoining, is emerging as strong and independent prognostic factor across multiple malignancies^1-7^. Reliable detection of chromothripsis requires whole genome sequencing (WGS) and the integration of both structural variants (SVs) and copy number (CN) data^1,2,4,8^.

Recently, we reported a comprehensive study of structural variation (SV) in a series of 752 newly diagnosed multiple myeloma (NDMM) from the CoMMpass trial for which long-insert low-coverage WGS was available (NCT01454297)^9^. Using the latest criteria for chromothripsis^1-5^ and manual curation, we reported a 24% prevalence of chromothripsis, making multiple myeloma (MM) the hematological cancer with the highest documented prevalence of chromothripsis^1,5,10,11^. In MM, chromothripsis has different features to that seen in solid cancers. Although the biological impact is likely similar across various malignancies, in MM and in other hematological malignancies, the structural complexity of each chromothripsis event is typically lower than seen in the solid cancers^1,4,5^. Specifically, the total focal CN gains within the regions of chromothripsis is lower than in solid organ cancer and in MM there is a lack of enrichment for double-minutes and other more catastrophic events such as *typhonas*^1,8^.

CN signatures have been reported in ovarian cancer as a potential BRCAness surrogate^12^. This important marker, denoting both prognosis and treatment-responsiveness, is detectable only by combining multiple WGS features^12,13^. Similarly, given the genome-wide distribution and complexity of chromothripsis in MM, we hypothesized that a comprehensive signature analysis approach using CN may provide an accurate estimation of chromothripsis in MM, without requiring specific SV assessment. Using the CoMMpass trial low-coverage long-insert WGS (n=752) and an additional validation set of WGS from NDMM (n=34) and other hematological malignancies (n=235), we demonstrate the accuracy and reproducibility of CN signatures for the detection of chromothripsis. In the CoMMpass dataset we show that CN signatures independently associate with shorter progression free (PFS) and overall survival (OS). Finally, to accelerate the clinical translation of testing for chromothripsis where WGS data is not available, we extended the analysis to whole exome sequencing (WES), where we confirm the ability of CN signatures to predict the presence of chromothripsis and show it is associated with adverse clinical outcomes.

## Results

### Experimental data and design

Genome-wide somatic CN profiles were generated from 752 NDMM patients with low-coverage long-insert WGS (median 4-8x) from the CoMMpass study (NCT01454297; IA13; **Supplementary Table 1**)^14,15^. The final SV catalog was generated by combining the two SV calling algorithms, DELLY^16^ and Manta^17^ with CN data, followed by a series of quality filters (see **Methods**)^9^. According to the most recently published criteria^1-5^, at least one chromothripsis event was observed in 24% of the entire series^9^. Patients with chromothripsis events were characterized by poor clinical outcomes, with chromothripsis being associated with multiple unfavorable clinical and genomic prognostic factors including translocations involving *MAF*, *MAFB* and *MMSET*, increased APOBEC mutational activity, del17p13 and *TP53* mutations^9^.

### De novo CN signature extraction in multiple myeloma

CN signature analysis takes the genome-wide CN gains and losses (**Figure 1a**), and measures 6 fundamental CN features: (i) number of breakpoints per 10 Mb, (ii) absolute CN of segments, (iii) difference in CN between adjacent segments, (iv) number of breakpoints per chromosome arm, (v) lengths of oscillating CN segment chains, and (vi) the size of segments (**Figure 1b**)^12^. The optimal number of categories in each CN feature was established using a mixed effect model with the *mclust* R package (**Figure 1c-d**). The consequence of taking this approach is that different malignancies and types of sequencing data may result in varying numbers of CN categories and thresholds defining these categories (see **Methods**)^12^.

**Figure 1.**
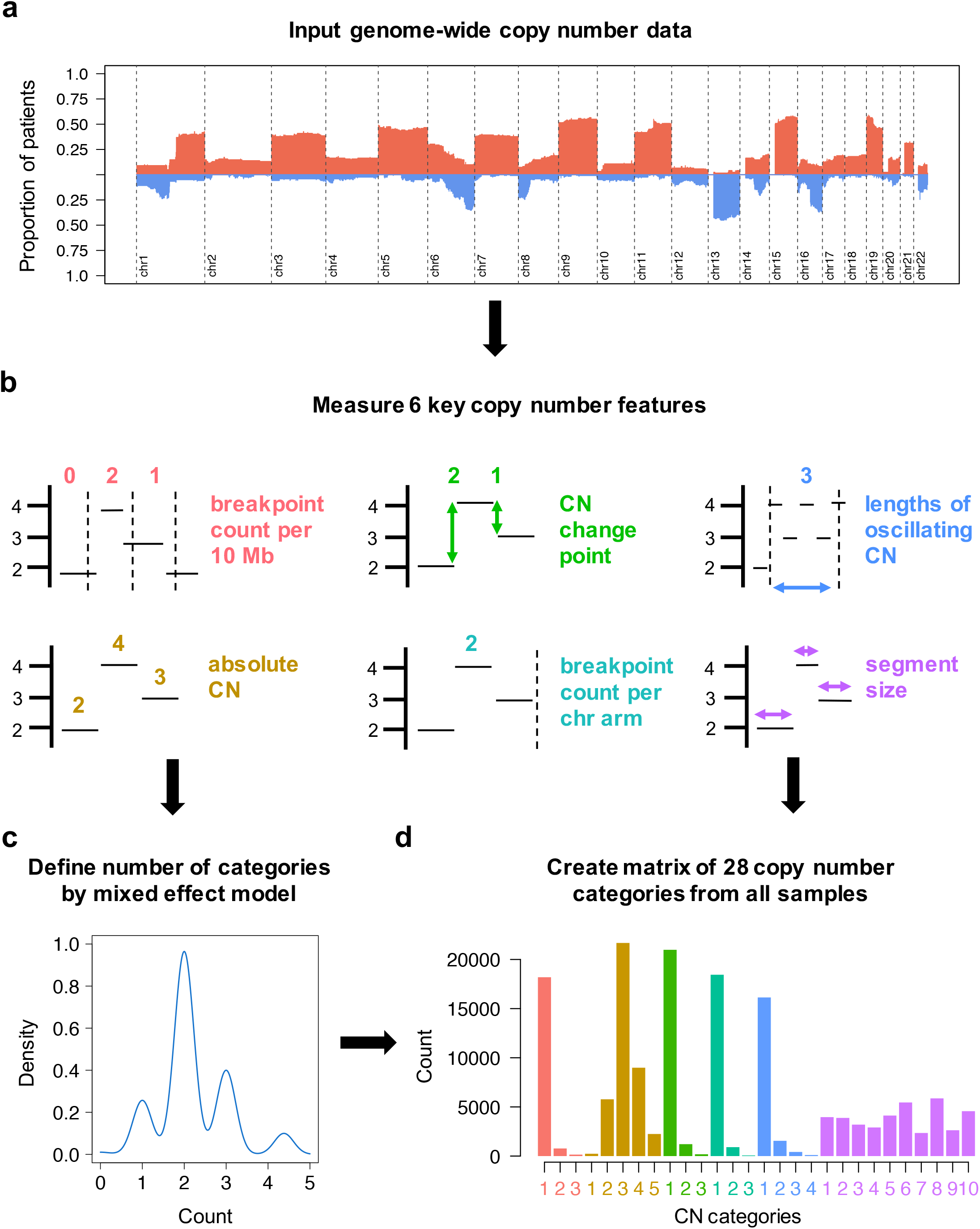
A schema demonstrating the definition of copy-number (CN) features from multiple myeloma whole genome sequencing data. **a**) Input genome-wide copy number gain and loss data from 752 newly diagnosed multiple myeloma whole genomes. **b**) Measure copy number as classified by 6 key features. **c**) Define the optimum number of categories for each copy number feature by a mixed-effects model (*mclust*). **d**) Tally the number of CN variation for each of 28 CN categories to produce a matrix of key CN features. This comprises the input matrix for the hierarchical Dirichlet process (*hdp*) for *de novo* extraction of CN signatures.

To take account of the biology of MM, we introduced a few modifications to the original CN features described by Macintyre et. al.^12^: (i) given the known poor quality mapping and copy number complexity related to class switch recombination and VDJ rearrangements, the regions corresponding to IgH, IgL and IgK were removed; (ii) considering both the low-coverage long-insert WGS limitation for calling subclonal copy number events and the less complex MM karyotype compared to solid cancers, fixed criteria for copy number status were introduced (see **Methods**, and **Supplementary Data 1** for full analytical R code).

Analyzing the CoMMpass long-insert low-coverage WGS; 28 CN categories were defined (**Figure 1d**; **Supplementary Table 2**). In comparison to the CN features described in ovarian cancer, in MM we observe lower total CN (median; 2, maximum; 9, compared with total CN exceeding 30 in a proportion of ovarian cancer)^12^. We also note shorter lengths of oscillating CN, and a low contribution from very large aberrant segments (in comparison to the dominant contribution from segments >30Mb in ovarian CN signature #1)^12^. Overall, these differences are in line with the lower genetic complexity of MM compared to ovarian cancer^18^.

Running the hierarchical Dirichlet process (*hdp*), 5 CN signatures were extracted in MM (**Figure 2**; **Supplementary Table 3**). CN-SIG1, CN-SIG2 and CN-SIG3 have high contributions from CN categories representing low numbers of breakpoints per 10Mb and breakpoints per chromosome arm. These signatures have small absolute differences between adjacent CN segments and short lengths of oscillating copy number. Each signature varies in the distribution of segment size and in the relative contribution of each CN category; CN-SIG1 has minimal jumps between adjacent segments and a higher contribution from larger segment sizes, mostly single chromosomal gains and trisomies. CN-SIG2 has higher total CN (i.e. multiple chromosomal gains and tetrasomies) and a higher contribution from small segments without jumps between adjacent segments; and CN-SIG3 is enriched for low absolute CN (i.e. deletions) with usually isolated events (rare oscillating events or multiple events on the same arm/chromosome) (**Figure 2**). In contrast, CN-SIG4 and CN-SIG5 were characterized by higher numbers of breakpoints per 10Mb and per chromosome arm, longer lengths of oscillating CN, and a higher contribution from small segments of CN change. While, CN-SIG4 has contribution from each of the 3 categories reflecting longer oscillation lengths, CN-SIG5 was characterized by a higher contribution from jumps in CN between adjacent segments, and by a higher contribution from high absolute CN (**Figure 2**).

**Figure 2.**
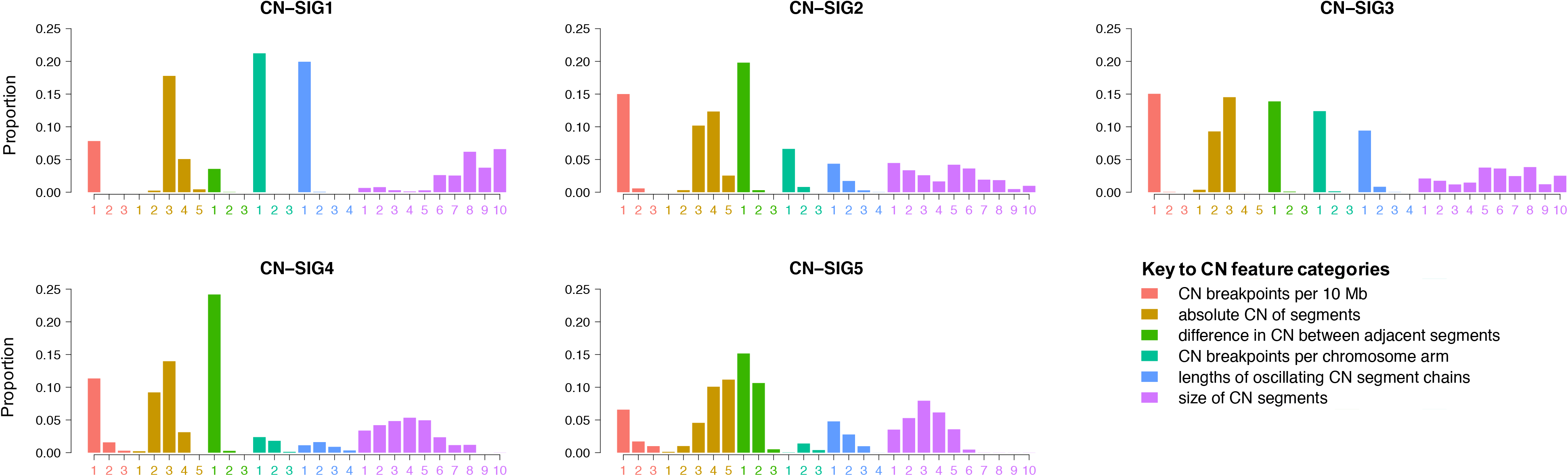
*De novo* extraction from whole genome sequencing data produces 5 copy-number (CN) signatures in 752 newly diagnosed multiple myeloma. The 5 CN signatures extracted comprise varying contribution across the 28-CN-feature matrix. The 2 chromothripsis-associated signatures are CN-SIG4 and CN-SIG5. (CN-SIG: copy-number signature).

### CN signatures are strongly predictive of chromothripsis in multiple myeloma

Given the complex CN features noted in CN-SIG4 and CN-SIG5 (**Figure 2**), we examined the association of these signatures with known MM genomic features^9,14,19-22^. Both signatures were correlated with features of high-risk MM (**Figure 3a**), including translocations involving *MAF*/*MAFB* (p=0.0005), APOBEC mutational activity (i.e. mutational signatures^23,24^; p<0.0001), biallelic *TP53* inactivation (p<0.0001), and 1q21 gain/amplification (p<0.0001; **Figure 3b-e**). There was a negative association with t(11;14)(*CCDN1*;*IGH*) (p<0.0001; **Supplementary Figure 1a**), consistent with the relative genomic stability known to be associated with a large proportion of this molecular subgroup of MM^14,19^. We show that CN-SIG4 and CN-SIG5 are highly correlated with the presence of complex structural chromosomal rearrangements (**Figure 3a**), including the subgroup of chromoplexy (p<0.0001; **Figure 3f**), and rearrangements defined as “complex- not otherwise specified” (complex-NOS; p<0.0001; **Supplementary Figure 1b**; **Methods**). Interestingly, the largest significant difference was noted with chromothripsis; with a median contribution of CN-SIG4/5 of 0.33 being seen in those cases with chromothripsis [inter-quartile range (IQR) 0.20-0.48] compared with 0.05 in those without (IQR 0.02-0.13) (p<0.0001; **Figure 3g**).

**Figure 3.**
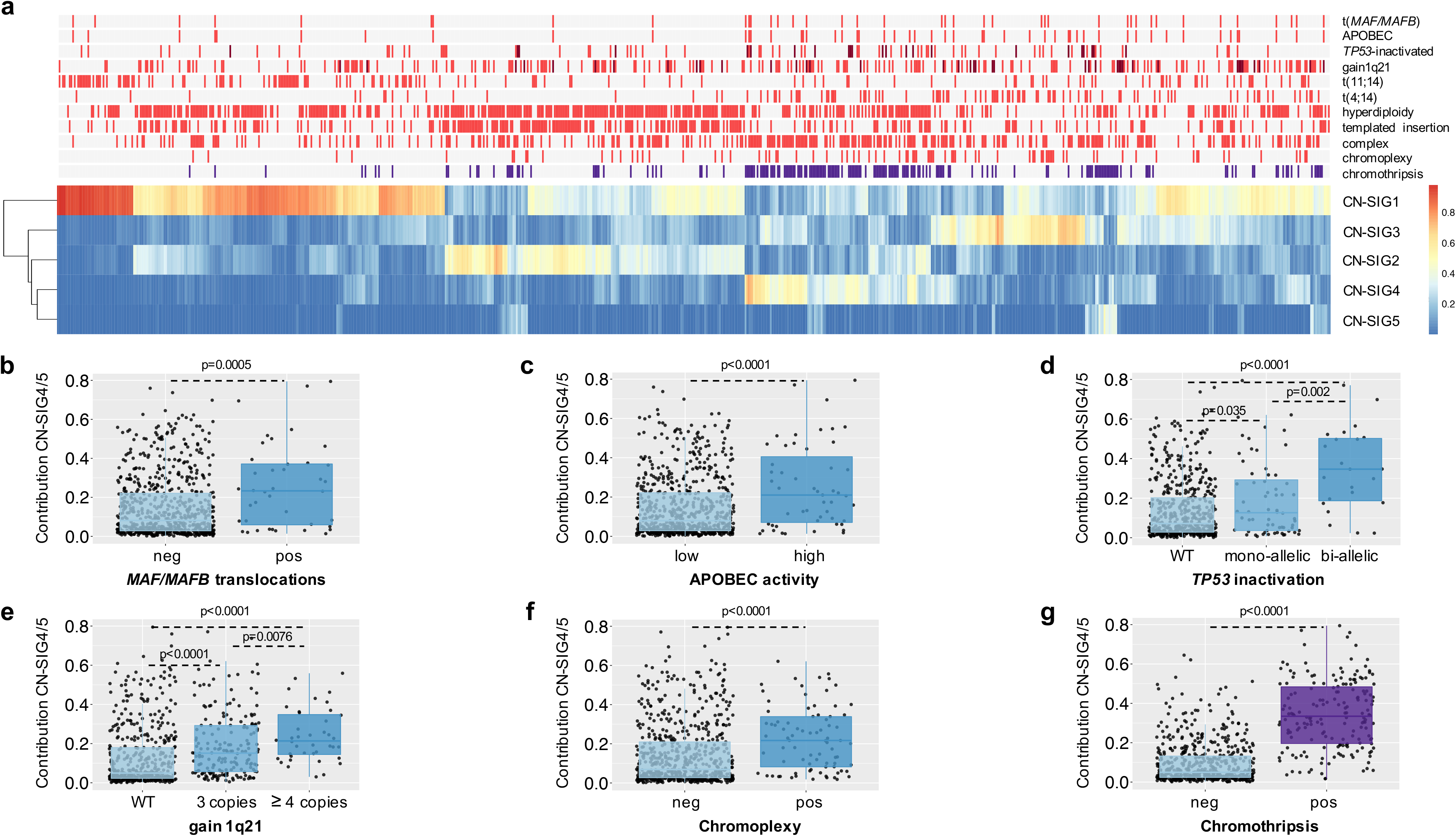
Clinical data demonstrates the correlation of copy number (CN) signatures with high-risk multiple myeloma prognostic features and complex genomic change. **a**) A heatmap of MM mutational and structural features demonstrates that contribution from CN-SIG4 and CN-SIG5 cluster with features of high-risk MM. Presence of biallelic *TP53* inactivation and chromosome 1q21 amplification (i.e. >3 copies) are annotated in dark red; presence of chromothripsis in purple; all the other genomic features are in bright red when present. **b-g**) There is a significantly higher median contribution from CN-SIG4 and/or CN-SIG5 on the samples having translocations involving (**b**) *MAF*/*MAFB*, (**c**) increased APOBEC mutational activity, (**d**) biallelic *TP53* inactivation, (**e**) 1q21 gain/amplification, (**f**) chromoplexy and (**g)** chromothripsis. Each boxplot shows the median and inter-quartile range (IQR) contribution of CN-SIG4 and CN-SIG5 across all patients, with whiskers extending to 1.5 * IQR. (Neg; lacking the feature; pos: containing the feature, WT: wild type).

These data suggest that CN signature analysis has the potential to accurately predict the presence of chromothripsis from WGS data derived from MM patients (**Figure 4a-b**). Evaluation of prediction accuracy of CN signatures by receiver operating curve (ROC) analysis with 10-fold cross validation confirmed this hypothesis, showing that CN signatures are highly predictive of chromothripsis; producing an average area-under-the-curve (AUC) of 0.90 (**Figure 4c**; **Supplementary Table 4**; for the full analytical R code, see **Supplementary Data 2**).

**Figure 4.**
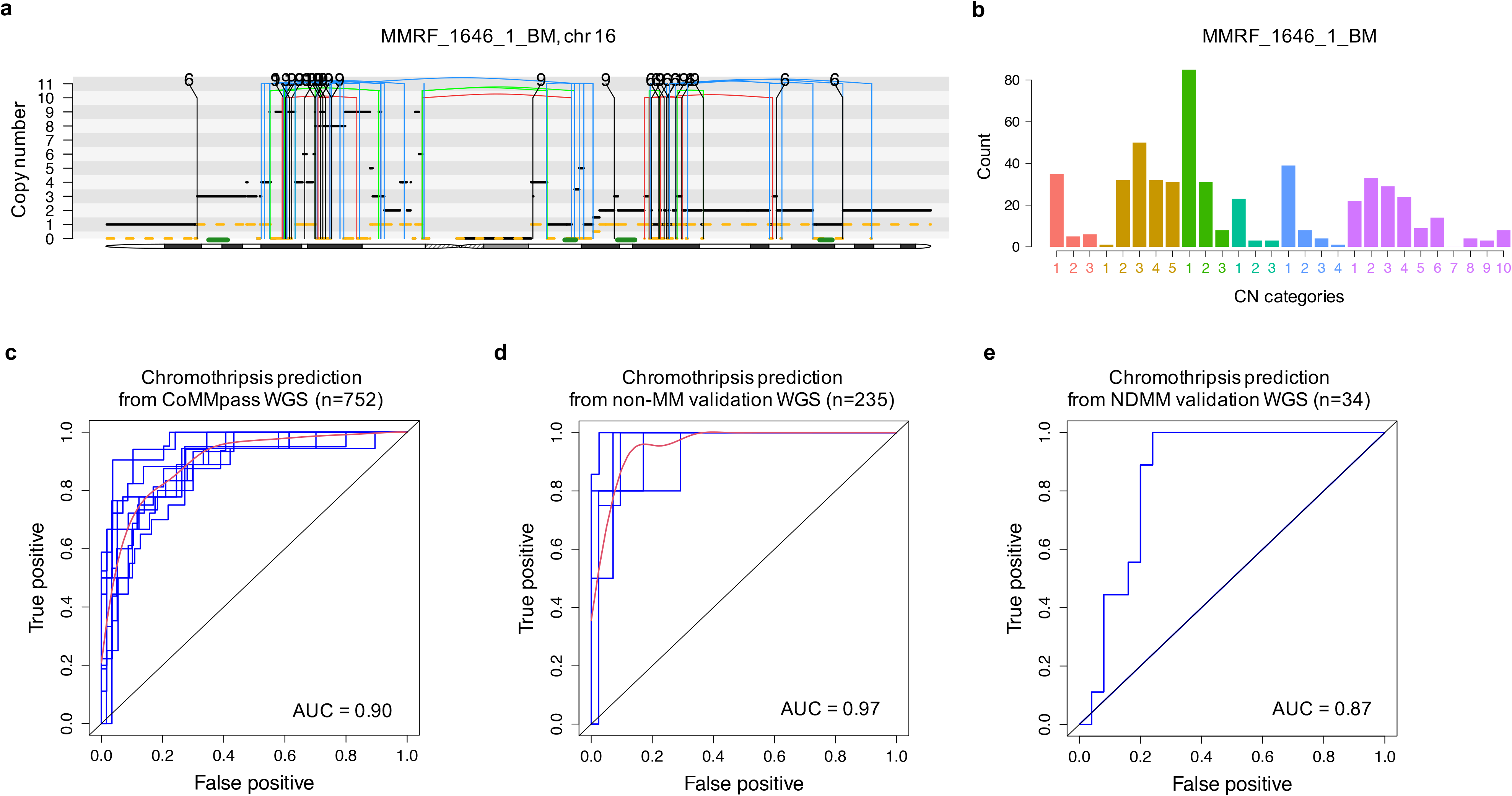
Copy number (CN) signatures in newly diagnosed multiple myeloma are strongly predictive of chromothripsis. **a**) An example of chromothripsis from the CoMMpass dataset (MMRF_1646_1_BM; chr: chromosome). The horizontal black line indicates total copy number; the dashed orange line minor copy number. Vertical lines represent SV breakpoints for deletion (red), inversion (blue), tandem-duplication (green) and translocations (black). **b**) the CN category profile from the same example patient (MMRF_1646_1_BM). **c**) Receiver operating curve (ROC) for the prediction of chromothripsis from CN signature analysis of CoMMpass whole genome sequencing (WGS) data (n=752). **d**) ROC for the prediction of chromothripsis from the validation set of other hematological cancers (n=235). For **c-d**: blue lines represent individual ROC (from 10-fold cross validation in **c** and 5-fold validation in **d**), red lines represent the mean of individual ROC, AUC: mean area-under-the-curve. **e**) ROC for the prediction of chromothripsis from the newly diagnosed multiple myeloma subset of the validation WGS (n=34).

### CN signatures are strongly predictive of chromothripsis in hematological malignancies

Given the low documented prevalence and low complexity of chromothripsis in hematological cancers^1,10,11^, we validated our prediction model using an extended dataset of 269 full coverage WGS from previously published hematological cancer samples, including data from the Pan-Cancer Analysis of Whole Genomes (PCAWG) study (n=269) ^5,25,26^. This included 34 NDMM, 92 chronic lymphocytic leukemia, 29 chronic myeloid leukemia, 104 B-cell lymphoma and 10 acute myeloid leukemia (7 *de novo*, 3 therapy-related) (**Supplementary Table 5**; **Methods**). Overall, the number of categories extracted in this series of WGS was smaller compared to the CoMMpass cohort (26 vs 28), likely reflecting the less impaired cytogenetic profile of non-MM hematological cancers^4^. Following the same computational approach reported in **Supplementary Data 1** and **2**, *de novo* extraction on the entire validation cohort identified 4 CN-signatures which were highly similar to those described in the CoMMpass WGS (**Supplementary Figure 2**; **Supplementary Tables 6-7**). Across the cohort of non-MM hematological malignancies, (n=235), the resultant ROC analysis had an average AUC of 0.97 for predicting chromothripsis (**Figure 4d**, using 5-fold cross validation due to the smaller sample size), while an AUC of 0.87 was observed when testing only in NDMM (n=34) (**Figure 4e**). These data demonstrate the reproducibility of chromothripsis prediction from CN signatures, in both a separate set of hematological cancer WGS, and an independent set of NDMM samples.

### CN signatures are strongly predictive of clinical outcomes in multiple myeloma

Survival analysis on the CoMMpass data demonstrates that the presence of chromothripsis predicts for a shorter PFS and OS compared with those without^9^; median PFS of 32.2 months (95% confidence interval [CI] 25.2-48.3m) in those harboring chromothripsis compared with 41.1m (95%CI 37.8-47.2m) in those without (p=0.00011; **Supplementary Figure 3a**), and median OS of 53.3m with chromothripsis but not reached [NR] in those without (p<0.0001; **Supplementary Figure 3b**). Survival probability according to the CN-signature predictive model mirrored survival according to chromothripsis, with a median PFS of 29.7m (95%CI 25.2m- NR) in those demonstrating a high CN-signature prediction score (CN_pred) compared with 41.8m (95%CI 38.0-48.1m) in those with a low score (p=0.0017; **Figure 5a**; **Supplementary Table 4**; **Methods**). Median OS in those with high CN_pred score was also significantly shorter at 53.1m compared with NR in those with a low score (p<0.0001; **Figure 5b**).

**Figure 5.**
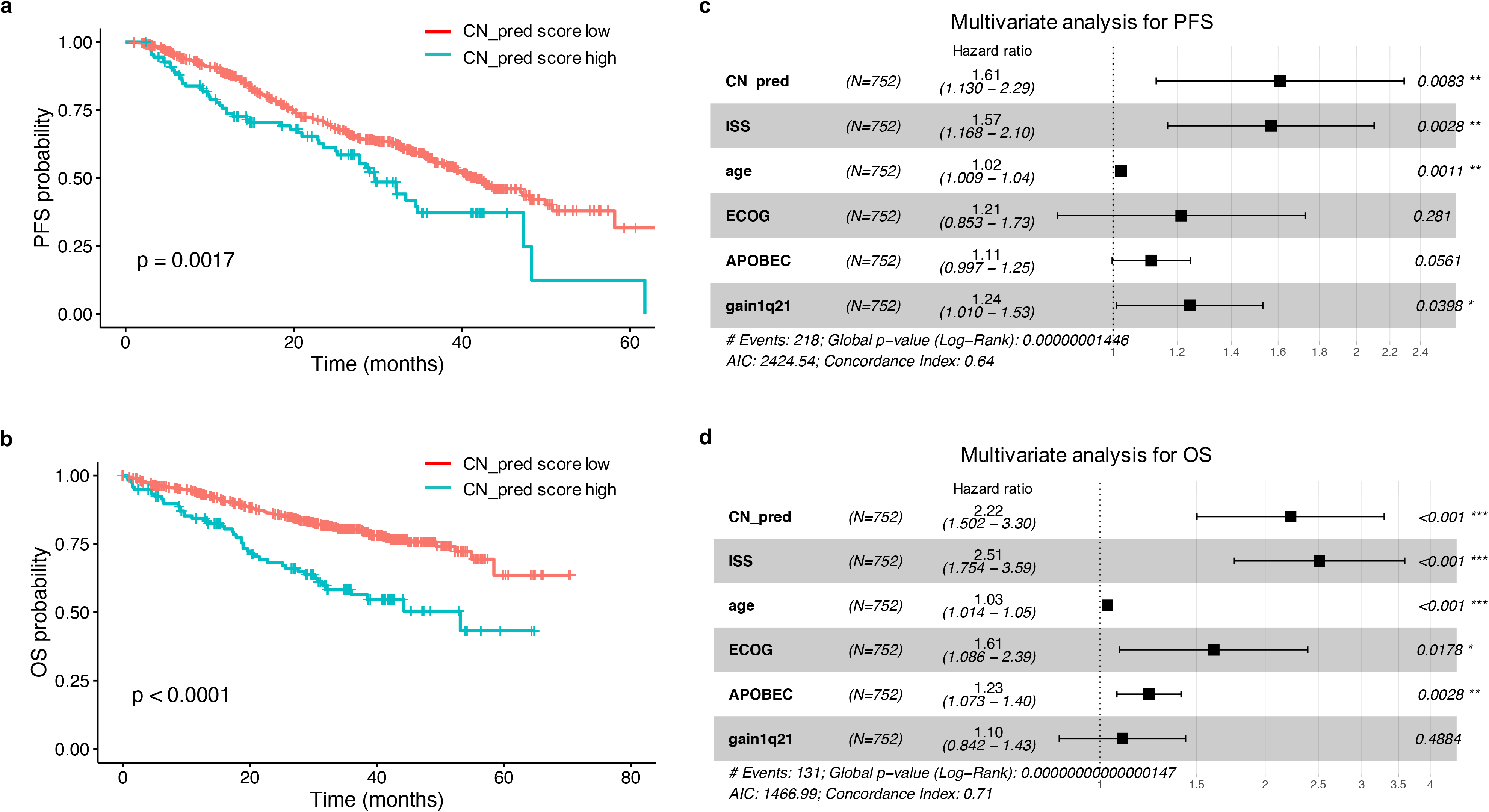
Copy number (CN) signatures in newly diagnosed multiple myeloma are independently predictive of clinical outcomes. **a**) Progression-free survival (PFS) probability in the CoMMpass dataset according to high (blue) or low (red) CN-prediction score for chromothripsis (CN_pred). **b**) Overall survival (OS) probability in the CoMMpass dataset according to high (blue) or low (red) CN_pred. **c**) Multivariate analysis of the effect of CN_pred on PFS after correction for International Staging Score (ISS), age, Eastern Cooperative Oncology Group (ECOG) score, 1q21 gain/amplification, and APOBEC mutational activity. **d**) Multivariate analysis of the effect of CN_pred on OS after correction for the same factors.

To select most important features from highly correlated genomic risk factors (**Figure 3a**) we performed a backwards stepwise cox regression including ISS, age, ECOG status, biallelic *TP53* inactivation, t(4;14)(*FGFR3*;*IGH*), 1q21 gain/amplification, increased APOBEC mutational activity and *MAF*/*MAFB* translocations. Based on this approach the final model contained ISS, age, ECOG, APOBEC mutational activity, 1q21 gain/amplification, and the CN_pred score. The model is consistent with previous published data indicating that APOBEC mutational activity is one of the strongest adverse prognostic factors in MM^20,22,27^, and that 1q21 gain/amplification is associated with early relapse^28^. The CN_pred score showed a significant association with shorter PFS and OS after controlling for other variables in the model, producing a hazard ratio (HR) of 1.61 (95% CI 1.13-2.29, p=0.0083, **Figure 5c**), and 2.22 (95% CI 1.50-3.30, p<0.001, **Figure 5d**), respectively.

### CN signatures compared with other CN-based tools

We next compared the prediction of the presence of chromothripsis by CN signatures with other CN-based algorithms recently used in MM to identify high-risk disease: a loss-of-heterozygosity index (LOH_index)^14,29^ and the genomic scar score (GSS)^27,30^ (see **Methods**). Results from each of these CN assessment approaches showed a right-skewed distribution across the CoMMpass 752 NDMM, [LOH_index; median 2, (range 0-27)**, Supplementary Figure 4a**, and GSS; median 7, (range 0-39), **Supplementary Figure 4b**], with the GSS distribution closely resembling that of previously published data in NDMM^27^.

Each of the LOH_index and the GSS demonstrated a lower average AUC for predicting the presence of chromothripsis in MM WGS (0.69 and 0.78 respectively, **Supplementary Figure 4c-d**). The difference in chromothripsis prediction between CN signatures and the LOH_index is quantitated as a statistically significant difference of 0.21 in AUC [based on bootstrap analysis, standard deviation(SD)=0.006, p<0.0001, **Supplementary Figure 4e**] while the difference in prediction between CN signatures and the GSS is quantitated as a statistically significant difference of 0.13 in AUC (based on bootstrap analysis, SD=0.005, p<0.0001, **Supplementary Figure 4f**).

In order to compare the effect of these different CN models on PFS and OS in multivariate analysis, the CN-signature prediction data was used as linear variable, which after correction for the previously included risk factors was associated with shorter PFS (HR=1.87, 95%CI 1.16-3.01, p=0.0106; **Supplementary Figure 5a**) and OS (HR=3.1, 95%CI 1.84-5.4, p<0.001, **Supplementary Figure 5b**). Performing multivariate analysis for PFS with correction for the same risk factors showed that neither the LOH_index (PFS HR=1.03, 95% CI 0.99-1.08, p=0.19; **Supplementary Figure 5c**) nor the GSS (PFS HR=1.02, 95% CI 1.0-1.04, p=0.12; **Supplementary Figure 5e**) retain a significant association. Each model has a slightly increased HR for OS in multivariate analysis; (LOH_index HR=1.1, 95% CI 1.02-1.1, p=0.008, **Supplementary Figure 5d**; GSS HR=1.0, 95% CI 1.02-1.1, p=0.001, **Supplementary Figure 5f**). Overall, CN-signatures perform significantly better at predicting poor outcomes in comparison with either the LOH_index or the GSS, suggesting that a more accurate prediction of chromothripsis is a better tool for identifying prognosis using CN-based information.

### CN signatures predict chromothripsis and clinical outcomes in whole exome sequencing data

Any prognostic assessment for MM would ideally be applicable in non-WGS assays, as WGS is currently both expensive and computationally intensive, making its clinical application outside of a research setting difficult. We performed *de novo* signature extraction using WES data from 677 NDMM CoMMpass samples, all of which also had WGS. The presence of these data enabled us to compare results in WES with the gold-standard method for chromothripsis-detection on WGS. The CN feature profile extracted from WES data was highly analogous to that obtained from WGS data (cosine similarity=0.99 for corresponding matrix columns), with a smaller contribution from the oscillation CN categories due to the lower data resolution overall, and in particular of focal and small lesions (**Supplementary Figure 6a**). *De novo* extraction using *hdp* produced 5 exome-based CN signatures (eCN), similar in their CN feature distribution to the signatures defined in WGS (**Supplementary Figures 6b-c**; **Supplementary Tables 7-8**). ROC analysis based on 10-fold validation produced an average AUC of 0.82 for predicting chromothripsis (**Supplementary Figure 7**; **Supplementary Table 4**).

The exome CN signature-based chromothripsis prediction score (eCN_pred) was associated with a significantly shorter PFS; median 26.0m (95%CI 18.0-48.3m) in those with a high eCN_pred score compared with 41.1m (95%CI 36.7-50.0m) in those with a low score, (p=0.0031; **Figure 6a**). OS was also significantly shorter; median 52.3m with a high eCN_pred score but NR in those with a low score, (p<0.0001; **Figure 6b**).

**Figure 6.**
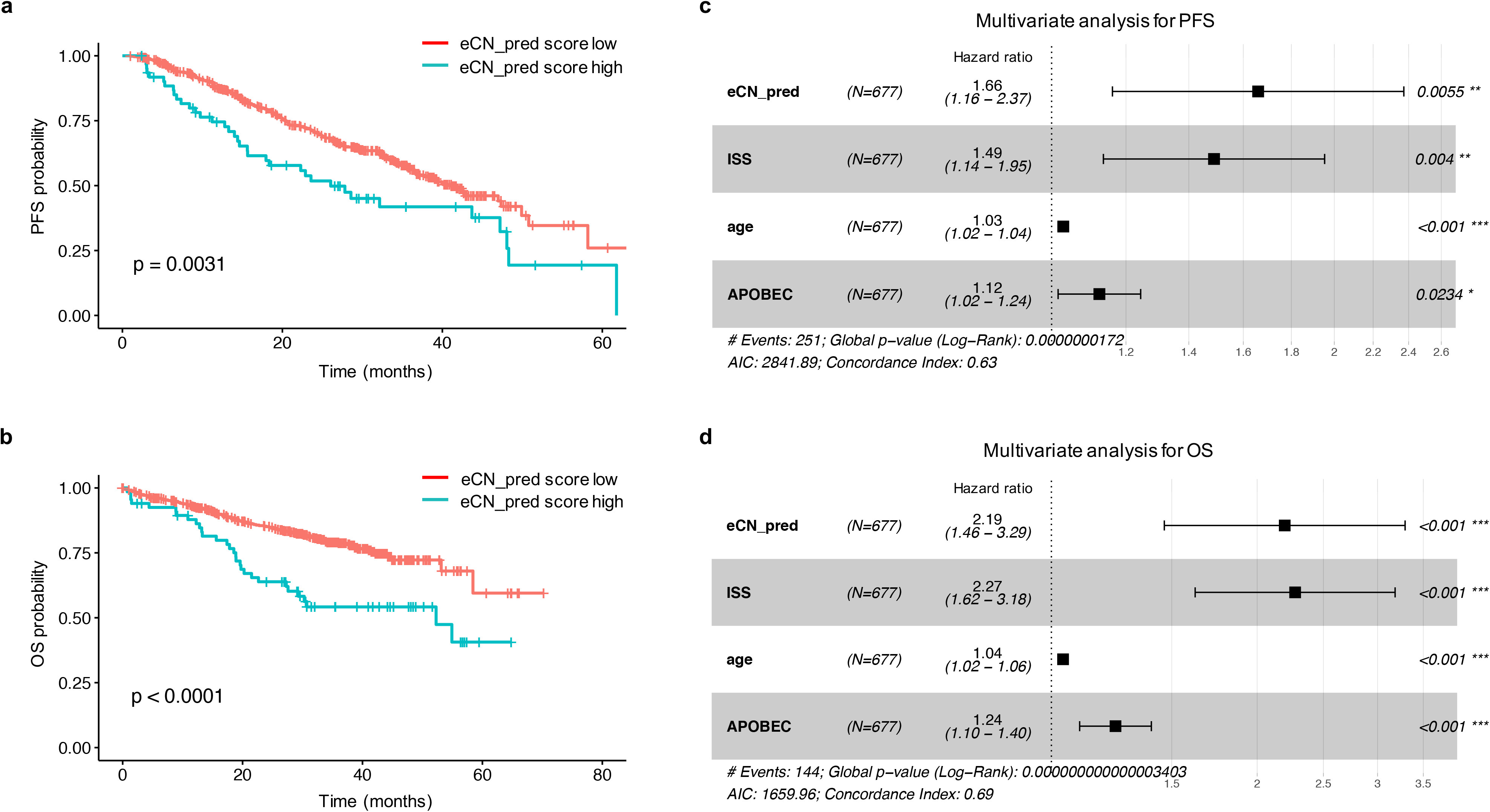
Copy number (CN) signatures extracted from whole exome sequencing (WES) in newly diagnosed multiple myeloma are highly predictive of clinical outcomes. **a**) Progression-free survival (PFS) probability in the CoMMpass dataset according to high (blue) or low (red) exome CN-prediction score (eCN_pred) for chromothripsis. **b**) Overall survival (OS) probability in the CoMMpass dataset according to high (blue) or low (red) eCN_pred. **c**) Multivariate analysis of the effect of eCN_pred on PFS after correction for International Staging Score (ISS), age and APOBEC mutational activity. **d**) Multivariate analysis of the effect of eCN_pred on OS after correction for the same factors.

In the exome data, backwards stepwise regression demonstrated that the best model for predicting survival was that comprising age, ISS, APOBEC-activity and the eCN_pred score. Multivariate analysis again produced a significant and independent association of eCN_pred with a shorter PFS (HR=1.66, 95%CI 1.16-2.37, p=0.0055; **Figure 6c**), and shorter OS (HR=2.19, 95%CI 1.46-3.29, p<0.001; **Figure 6d**) recapitulating both the results obtained from WGS CN signature based chromothripsis prediction (**Figure 5**), and those obtained by manual data curation^9^.

## Discussion

We recently carried out a comprehensive analysis of the landscape of SVs in MM, showing their critical role in disease pathogenesis and confirming the importance of WGS for deciphering the genomic complexity of these events ^9,19,31^. We demonstrated a high prevalence of complex structural events such as chromothripsis in MM (24%) which is comparable to that detected in other malignancies; recent data across 38 cancer types from the PCAWG consortium described high-confidence calls for chromothripsis events occurring in 29% of all samples, and above 50% in several cancer types (melanoma, sarcoma, lung adenocarcinoma)^1^. Given the adverse prognostic association of chromothripsis with PFS and OS in MM and other malignancies^7,9,32,33^, that is independent of other known prognostic variables, it follows that the integration of complex SV data has the potential to improve the current prognostic scoring systems.

Current approaches for identifying these complex SVs require expense and time commitment because of the need for either the manual curation of WGS data or the use of computational tools requiring both CN and SV data to predict chromothripsis^1,8,9,34^. In order to circumvent these issues, we investigated CN signature approach for predicting the presence of chromothripsis. This approach was initially developed in ovarian cancer as a potential surrogate for predicting BRCA deficiency^12^. We have extended this approach for use in MM and show that a CN signature analysis of NDMM provides an accurate prediction of the presence of chromothripsis that outperforms other CN assessment algorithms^27,29^. The survival probability identified using a CN signature-based prediction of chromothripsis closely mimics PFS and OS curves observed with the presence / absence of chromothripsis^9^. Using a validation set of WGS containing multiple hematological malignancies, we provide proof-of-principle that CN signature analysis can predict for chromothripsis across different hematological cancer types and can, therefore, be used as surrogate for these variants to further address the role of chromothripsis in these blood cancers.

The primary objective was to test whether WGS-based CN signatures can reliably predict chromothripsis and its poor impact on clinical outcomes. Another critical aspect of this study was to expand our investigations using non-WGS (i.e. exome-based) data. In WES data, multivariate analysis revealed a significant association between CN signatures and shorter PFS and OS. Indeed, the clinical impact of CN signatures was similar in WGS and WES data. This is important from a translational perspective because it provides an easier pathway towards clinical application in the standard of care setting of NDMM patients.

In conclusion, CN signature analysis can accelerate our ongoing quest to accurately define high-risk MM, and to translate WGS-based prognostication into the clinic.

## Methods

### Samples

All the raw data used in this study are publicly available. Somatic CN profiles for the definition of CN signatures in MM were generated from 752 NDMM patients with low-coverage long-insert WGS (median 4-8x) from the CoMMpass study. The CoMMpass study is a prospective observational clinical trial (NCT01454297) with comprehensive genomic and transcriptomic characterization of NDMM patients, funded and managed by the Multiple Myeloma Research Foundation (MMRF)^35^. The study is ongoing, with data released regularly for research use via the MMRF research gateway, https://research.themmrf.org. In this study, we used Interim Analysis (IA) 13.

The validation dataset of hematological cancer WGS was compiled from several sources. Data from the Pan-Cancer Analysis of Whole Genomes (PCAWG) study^5,25,26^ was accessed via the data portal http://dcc.icgc.org/pcawg/, comprising 92 chronic lymphocytic leukemia, 29 chronic myeloid leukemia, 104 B-cell lymphoma and 7 acute myeloid leukemia. An additional 3 therapy-related AML were included, with the WGS data available from European Genome-phenome (EGA) under the accession code EGAD00001005028. Together, these samples formed the non-MM validation WGS set (n=235). The MM validation dataset (n=34) comprised 28 NDMM, 4 monoclonal gammopathy of uncertain significance (MGUS), 2 smoldering MM (SMM) and 1 plasma cell leukemia (PCL). It was compiled from 3 studies which can be accessed from European Genome-phenome (EGA) and the database of Genotypes and Phenotypes (dbGAP) with accession codes EGAD00001003309, EGAS00001004467 and phs000348.v2.p1.

WES from 677 NDMM patients were accessed from the CoMMpass study as above, with each patient having concurrent WGS available for comparison.

### CNV signature analysis

Genome-wide somatic copy number (CN) profiles were generated from 752 NDMM patients with long-insert low-coverage WGS available from the CoMMpass study. Paired-end reads were aligned to the human reference genome (HRCh37) using the Burrows Wheeler Aligner, BWA (v0.7.8) and CN variation and loss-of-heterozygosity events were identified using tCoNuT (https://github.com/tgen/tCoNuT), with verification performing using controlFREEC^14,15^. We minimized the inclusion of artefacts by removing all CN changes smaller than 50kB and excluding the regions corresponding to IgH, IgL and IgK, as well as the X chromosome from analysis.

The optimal number of categories in each of the 6 CN features detailed in **Figure 1** were established using a mixed effect model with the *mclust* R package, producing a CN category matrix with defined limits for each feature (**Supplementary Table 2**). Given the lower complexity of MM CN changes compared to the original CN signature definition in ovarian cancer^12^, fixed criteria for copy number status were introduced (#1 = bi-allelic deletion; #2 = monoallelic deletion; #3 = diploid; #4 = single gain; 5# = two or more gains, **Supplementary Data 1**). *De novo* CN signature extraction was performed from this matrix via the hierarchical Dirichlet process (*hdp*, https://github.com/nicolaroberts/hdp). The extracted CN signatures were then correlated with publicly available clinical data and manually curated SV data as detailed below to allow the calculation of prediction metrics.

The accuracy of chromothripsis prediction from CN signatures was assessed by the area-under-the-curve (AUC) from receiver operating characteristic (ROC) curves via 10-fold cross validation, using all extracted CN signatures as input. The sensitivity and specificity of chromothripsis prediction from varying levels of probability (i.e. AUC) were compared, (**Supplementary Table 4**), with a prediction level ≥ 0.6 defining a high CN_pred (WGS) and eCN_pred (WES) score. This score provided the highest Ievel of sensitivity for chromothripsis prediction while still keeping the specificity level at / above 95% for both WGS- and WES-based prediction.

Somatic variant calling was performed using DELLY (v0.7.6)^16^ and Manta (v.1.5.0)^17^. The final catalogue of high-confidence SVs was obtained by integrating DELLY and Manta calls with copy number data and applying a series of quality filters^9^. Briefly, all SVs called and passed by both callers were included and SVs called by a single caller were only included in specific circumstances: (i) SVs supporting copy-number junctions, (ii) reciprocal translocations, and (iii) translocations involving an immunoglobulin locus (i.e., *IGH, IGK*, or *IGL*).

SV single and complex events definition was according to the most recent consensus criteria^1-5^. Chromothripsis was defined by more than 10 interconnected SV breakpoint pairs associated with oscillating CN across one or more chromosomes; definition included: (i) clustering of breakpoints, (ii) randomness of DNA fragment joins, and (iii) randomness of DNA fragment order across one or more chromosomes. Chromoplexy was defined by interconnected SV breakpoints across >2 chromosomes associated with CN loss. Templated insertions were defined as a concatenation of translocations usually associated with focal CN gain; if >2 chromosomes were involved these events were classified as complex. Patterns of 3 or more interconnected breakpoint pairs that did not fall into the above categories were classified as “complex”, not otherwise specified^9^.

The majority of the clinical association data was obtained directly from the CoMMpass data portal (https://research.themmrf.org). The definition of high APOBEC activity was obtained from single-base substitution (SBS) signature analysis; a mutational signature fitting approach using the R package *mmsig*, (https://github.com/evenrus/mmsig) was applied to single nucleotide variant calls from WES data^25,26^. High APOBEC mutational activity was defined by an absolute contribution of APOBEC-associated signatures (SBS2 and SBS13) in the top decile, among patients with evidence of APOBEC activity^9,26^.

CN variation data from the validation dataset of hematological cancers was utilized for *de novo* CN signature extraction (hCN-SIG, **Supplementary Table 6**) without reference to the CoMMpass WGS-derived CN signatures. Fixed criteria for copy number status were introduced as detailed above. The presence of chromothripsis was confirmed by manual inspection of SV and CN data. The accuracy of chromothripsis prediction from was assessed by AUC from ROC curves using all extracted hCN signatures as input. 5-fold cross validation was used for the non-MM cohort prediction, which was then used as the training model for testing the prediction from the MM validation cohort.

CN variation and loss-of-heterozygosity events from the CoMMpass WES sequencing data was assessed using FACETS (Fraction and Allele specific Copy number Estimate from Tumor/normal Sequencing, https://github.com/mskcc/facets)^36^. Fixed criteria for copy number status were introduced as detailed above, then *de novo* CN signature extraction, and clinical / genomic correlation were all performed without reference to the WGS-derived CN signatures.

### Comparison with alternate CN assessment approaches

The LOH_index and the GSS were calculated from allele-specific CN files, with the methods being applicable to either WGS or WES data. The LOH_index was calculated using the R package *signature.tools.lib^29,37^* (https://github.com/Nik-Zainal-Group/signature.tools.lib), while the GSS was calculated using the R package *scarHRD*^30^ (https://github.com/sztup/scarHRD). The *scarHRD* output is 3 separate CN features (loss-of-heterozygosity, telomeric allelic imbalance, and number of large-scale transitions) which are summed to produce a final score.

To compare chromothripsis-prediction from CN signatures with each of the LOH_index and the GSS, we first calculated with difference in average AUC between two methods estimated from 10-fold cross-validation. Then, standard deviation of the difference in AUCs was estimated by performing a bootstrap resampling. On each new bootstrap sample, we estimated difference in the average AUC between two methods using 10-fold cross-validation. This procedure was repeated 1000 times (**Supplementary Data 3**).

### Software and statistics

Analysis was carried out in R version 3.6.1. The analytical workflow in R for the *de novo* extraction of CN signatures is provided in **Supplementary Data 1**, the code for predicting chromothripsis from CN signatures is detailed in **Supplementary Data 2** and the approach to comparing 2 methods for predicting chromothripsis is presented in **Supplementary Data 3**. Key software tools noted throughout the workflow (including *mclust, hdp, survminer*, *pROC, mmsig, signature.tools.lib,* and *scarHRD*) are publicly available. Unless otherwise specified, we used the Wilcoxon rank sum test to test for differences in continuous variables between two groups and Fisher’s exact test for 2×2 tables of categorical variables.

### Data availability

Sequencing files are available at the EGA and dbGaP archives under the following accession codes:

- EGAD00001003309 and phs000348.v2.p1 WGS: 24 NDMM and 1 high risk smoldering multiple myeloma patient
- phs000748.v1.p1: WES and low coverage/long insert WGS sequencing data from 752 NDMM patients (CoMMpass trial; IA 13)
- EGAS00001004467: WGS data from 3 MM, 1 SMM, 1 PCL and 4 MGUS patients
- EGAD00001005028 WGS data from 3 therapy related AML patients.
- Pan-Cancer Analysis of Whole Genomes (PCAWG) study^5,25,26^ was accessed via the data portal: https://dcc.icgc.org/

## Acknowledgements

This work is supported by the Memorial Sloan Kettering Cancer Center NCI Core Grant (P30 CA 008748), the Multiple Myeloma Research Foundation (MMRF), and the Perelman Family Foundation.

F.M. is supported by the American Society of Hematology, the International Myeloma Foundation and The Society of Memorial Sloan Kettering Cancer Center.

K.H.M. is supported by the Haematology Society of Australia and New Zealand New Investigator Scholarship, the Royal College of Pathologists of Australasia Mike and Carole Ralston Travelling Fellowship Award, the Royal Australasian College of Physicians Dr Helen Rarity McCreanor Travelling Fellowship and the Snowdome Foundation.

G.J.M is supported by The Leukemia Lymphoma Society.

NB is supported by the European Research Council under the European Union’s Horizon 2020 research and innovation programme (grant agreement No. 817997)

## Author contributions

F.M. designed and supervised the study, collected and analyzed data and wrote the paper; O.L. supervised the study, collected and analyzed data and wrote the paper; K.H.M. collected, analyzed and interpreted the data and wrote the paper; G.M. interpreted the data and wrote the paper; E.H.R., An.D., Z.B., V.Y. and B.D. collected and analyzed data. M.H., B.Z., E.B., P.B., N.B., Y.Z., A.Do., A.L. collected data.

