## Supplementary Figures and Tables for "Copy number signatures predict chromothripsis and associate with poor clinical outcomes in patients with newly diagnosed multiple myeloma"

Maclachlan et al.

### Supplementary Figure Legends

**Supplementary Figure 1. Correlation of copy number (CN) signatures with t(11;14) and complex not otherwise specified (NOS) structural variant events. a**) The contribution of CN-SIG4 / CN-SIG5 is higher in those samples lacking t(11;14)(*CCDN1*;*IGH*) than in samples containing this translocation. **b**) The contribution of CN-SIG4 / CN-SIG5 is higher in those samples containing complex SV NOS than in those without. Each boxplot shows the median and inter-quartile range (IQR) contribution of CN-SIG4 and CN-SIG5 across all patients, with whiskers extending to 1.5 * IQR. (Pos: containing the feature; neg: without the feature).

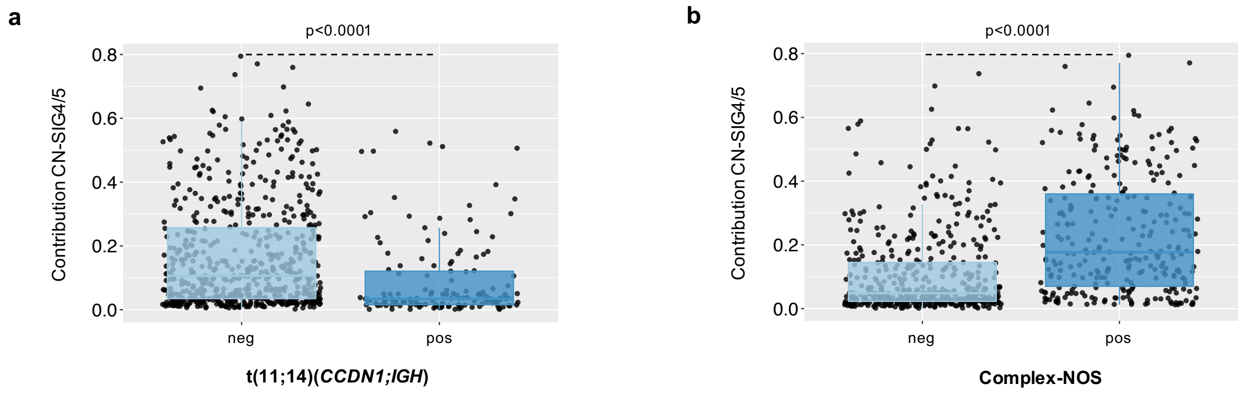

**Supplementary Figure 2. Copy number feature distribution from a validation set of whole genomes from hematological cancers. a)** Using *mclust* to define the optimum number of categories in each of the 6 copy number classes defined 26 categories in the validation set of WGS, which contains MM samples along with chronic lymphocytic leukemia, acute myeloid leukemia, and B-cell lymphoma. **b**) *De novo* extraction from the validation dataset extracted 4 CN signatures (defined as hCN-SIG to denote extraction from hematological cancer data). **c**) A heatmap of hCN-SIG in the validation WGS according to chromothripsis. (Purple: containing chromothripsis; grey: lacking chromothripsis).

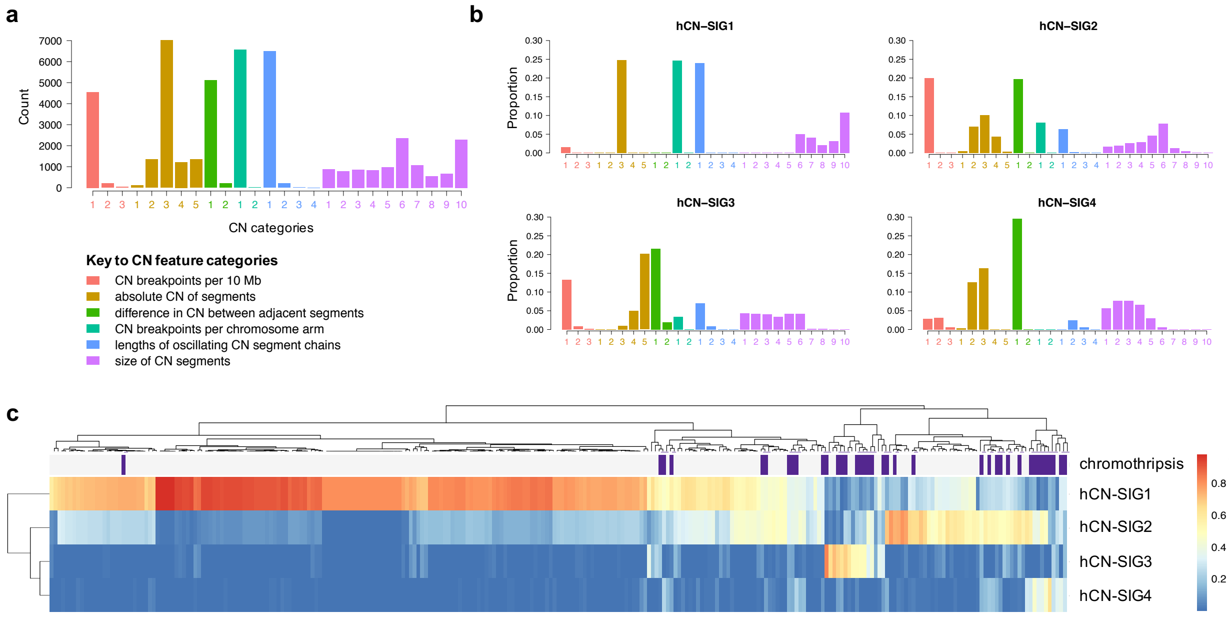

**Supplementary Figure 3. Survival according to copy number (CN) signatures in newly diagnosed multiple myeloma closely resembles survival according to the presence or absence of chromothripsis. a**) Progression-free survival (PFS) probability in the CoMMpass dataset according to the presence or absence of chromothripsis, and according to a high or low CN-prediction score for chromothripsis (CN_pred). **b**) Overall survival (OS) probability in the CoMMpass dataset according to the presence or absence of chromothripsis, and according to a high or low CN_pred score. (Chromothripsis; presence: green, absence: red; CN_pred; high: purple; low: blue).

**a**

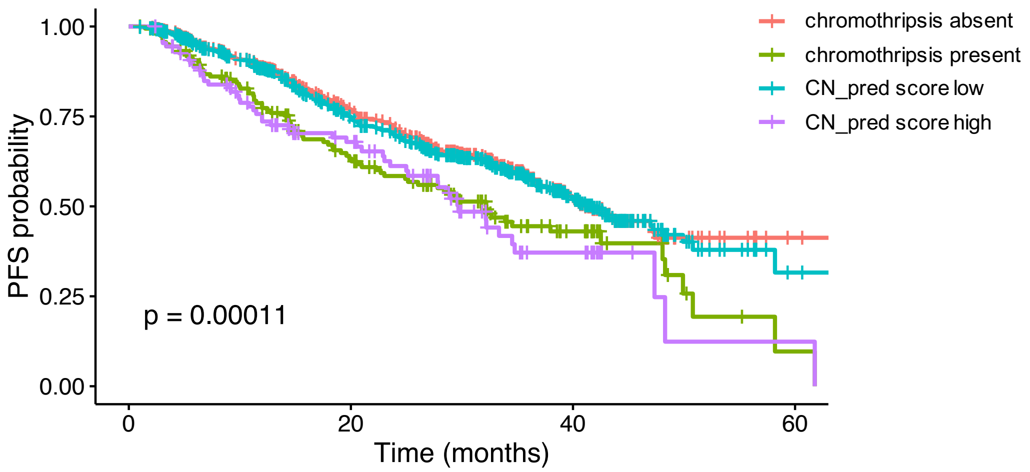


**b**

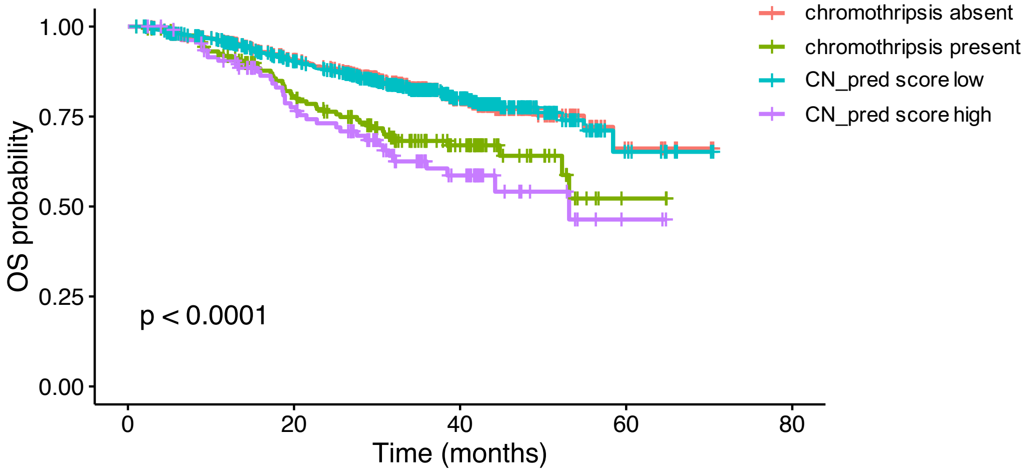

**Supplementary Figure 4. Comparison of the area-under-the-curve (AUC) estimates for the prediction of chromothripsis between copy number (CN) signatures, the loss-of-heterozygosity (LOH) index and the genomic scar score (GSS) in WGS. a-b**) Histograms presenting the distribution for each of (**a**) the LOH_index and (**b**) the GSS. **c-d**) Receiver operating curves (ROC) for the prediction of chromothripsis from the CoMMpass WGS data by (**c**) the LOH_index and (**d**) the GSS. **e-f**) Histograms presenting the results from bootstrap analysis comparing the difference in chromothripsis prediction between CN signatures and (**e**) the LOH_index and (**f**) the GSS.

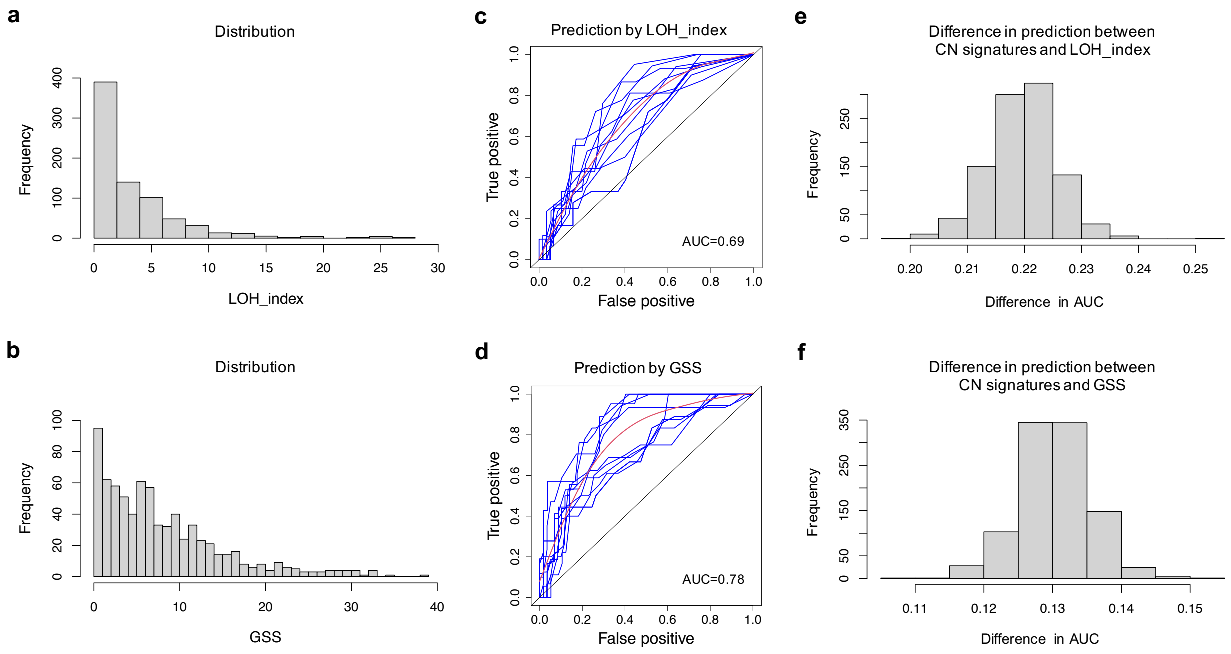

**Supplementary Figure 5. Comparison of progression free and overall survival between copy number (CN) signatures, the loss-of-heterozygosity (LOH) index and the genomic scar score (GSS) in whole genome sequencing.** Multivariate analysis of progression-free survival (PFS) from WGS after correction for age, International Staging Score (ISS), Eastern Cooperative Oncology Group (ECOG) score, APOBEC mutational activity and gain/amplification 1q21 according to (**a**) CN signature prediction as a linear feature (prediction), (**c**) the LOH_index and (**e**) the GSS. Multivariate analysis of the effect on overall survival (OS) of (**b**) CN signature prediction as a linear feature (prediction), (**d**) the LOH_index and (**f**) the GSS, after correction for the same factors.

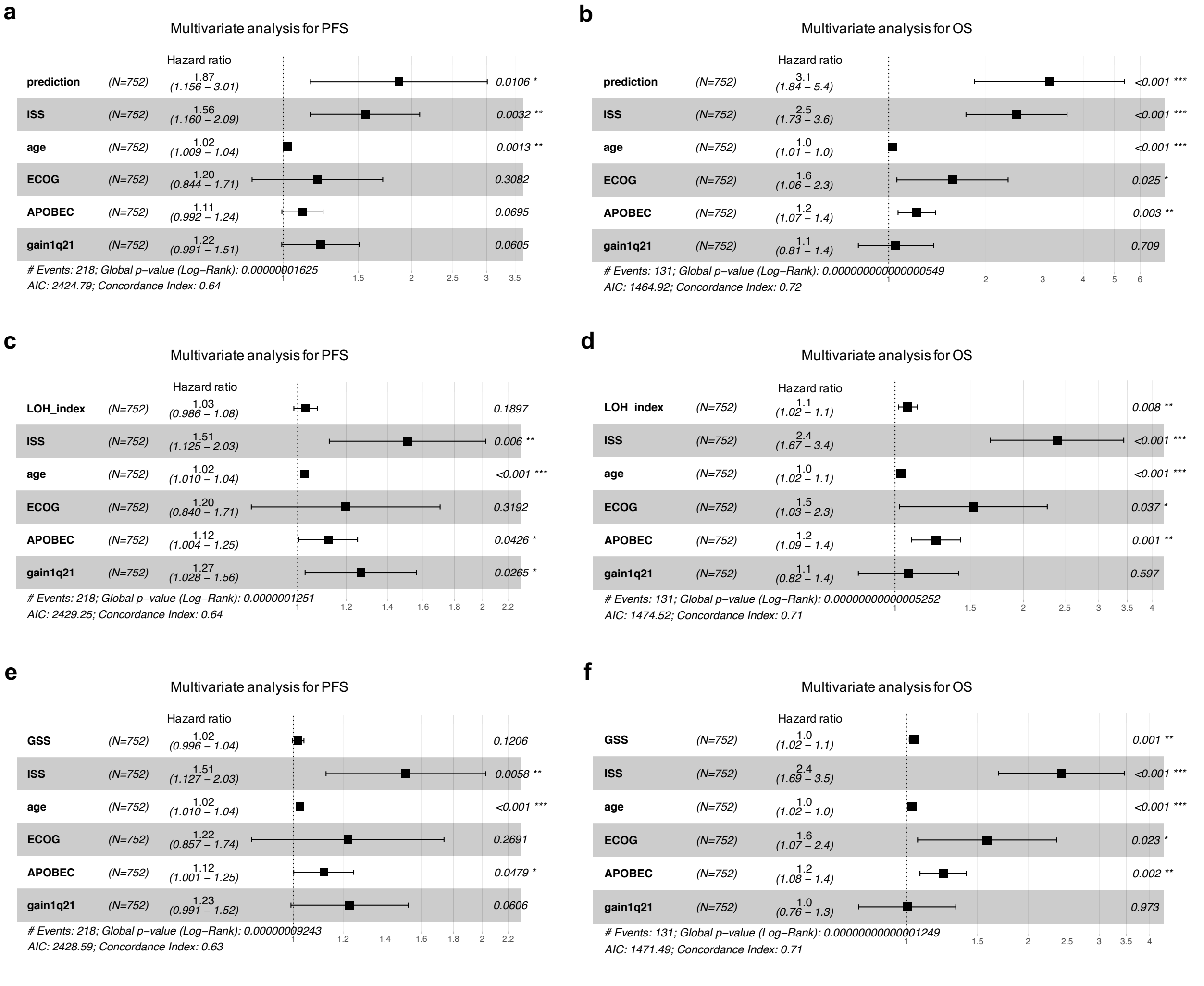

**Supplementary Figure 6. Copy number (CN) feature matrix and CN signatures from the CoMMpass whole exome sequencing (WES) data. a**) Using the CN feature limits defined in the CoMMpass WGS confirmed a similar CN feature matrix in the CoMMpass WES. **b**) *De novo* extraction without reference to WGS-derived CN signatures defined 5 CN signatures (defined as eCNV to denote extraction from exome data). **c)** A heatmap of eCN signatures from WES data clustered by APOBEC mutational activity, *MAF/MAFB* translocations, t(4;14), gain / amplification of chromosome 1q21, biallelic *TP53* inactivation and chromothripsis. Presence of biallelic *TP53* inactivation and chromosome 1q21 amplification (i.e. >3 copies) are annotated in dark red; presence of chromothripsis in purple; all the other genomic features are in bright red when present.

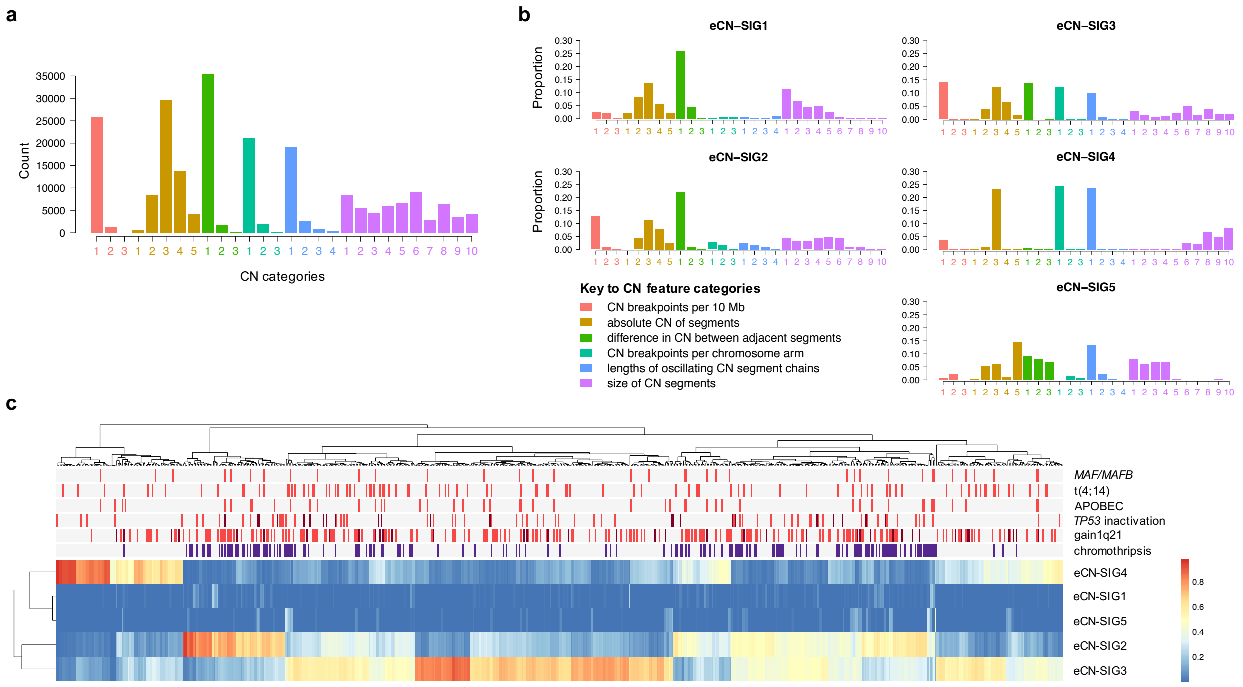

**Supplementary Figure 7. Copy number (CN) signatures extracted from whole exome sequencing (WES) in newly diagnosed multiple myeloma are highly predictive of chromothripsis.** Receiver operating curve (ROC) for the prediction of chromothripsis from CN signature analysis of CoMMpass WES data. (Blue lines: individual ROC from 10-fold cross validation; red lines: mean of individual ROC; AUC: mean area-under-the-curve).

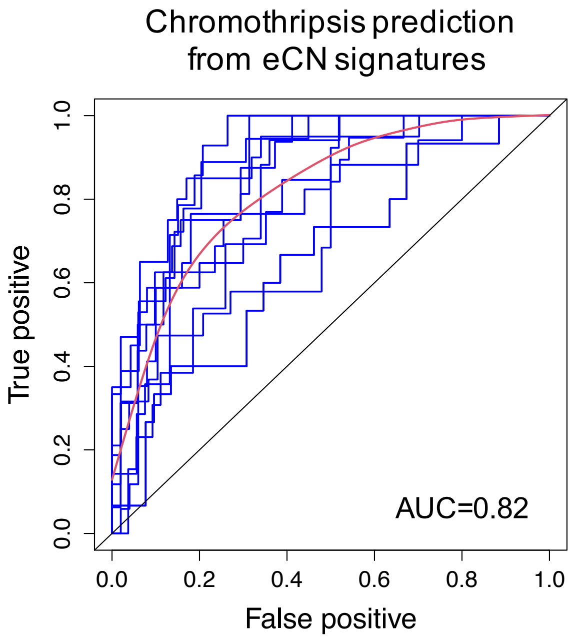

### Supplementary Table Legends

**Supplementary Table 1. Summary clinical details of patients from the CoMMpass dataset.**

| Clinical variable | Value |
|---|---|
| **Patients (n)** | 752 |
| **Gender (female, %)** | 39.9 |
| **Age (years, median, IQR)** | 64 (57 - 71) |
| **ISS stage III (%)** | 26.5 |
| **ECOG status ≥2 (%)** | 17.3 |
| **PFS follow-up (median, IQR, months)** | 22.7 (11.3-35.2) |
| **OS follow-up (median, IQR, months)** | 30.7 (15.9-41.0) |

ECOG = Eastern Cooperative Oncology Group; ISS = International Staging Score; IQR = interquartile range; n = number; PFS = progression-free survival; OS = overall survival.

**Supplementary Table 2. A matrix of 28 copy number (CN) categories across 6 CN features defined from newly diagnosed multiple myeloma patients in the CoMMpass dataset.**

| CN feature type | Category code | Count limit |
|---|---|---|
| breakpoints per 10 Mb | 1 | 3 |
| breakpoints per 10 Mb | 2 | 6 |
| breakpoints per 10 Mb | 3 | 31 |
| absolute CN | 1 | 0 |
| absolute CN | 2 | 1 |
| absolute CN | 3 | 2 |
| absolute CN | 4 | 3 |
| absolute CN | 5 | 9 |
| CN change point | 1 | 1 |
| CN change point | 2 | 3 |
| CN change point | 3 | 8 |
| breakpoints per chr arm | 1 | 5 |
| breakpoints per chr arm | 2 | 17 |
| breakpoints per chr arm | 3 | 60 |
| length of oscillating CN | 1 | 1 |
| length of oscillating CN | 2 | 4 |
| length of oscillating CN | 3 | 9 |
| length of oscillating CN | 4 | 38 |
| CN segment size | 1 | 165300 |
| CN segment size | 2 | 502740 |
| CN segment size | 3 | 1588400 |
| CN segment size | 4 | 7113100 |
| CN segment size | 5 | 21801700 |
| CN segment size | 6 | 53962800 |
| CN segment size | 7 | 67587881 |
| CN segment size | 8 | 105333496 |
| CN segment size | 9 | 142322700 |
| CN segment size | 10 | 249137896 |

**Supplementary Table 3. A matrix of copy number (CN) category contributions defining 5 CN signatures (CN-SIG) extracted from newly diagnosed multiple myeloma patients in the CoMMpass dataset.**

| CN feature type | Category code | CN-SIG1 | CN-SIG2 | CN-SIG3 | CN-SIG4 | CN-SIG5 |
|---|---|---|---|---|---|---|
| breakpoints per 10 Mb | 1 | 0.0782 | 0.1500 | 0.1504 | 0.1134 | 0.0658 |
| breakpoints per 10 Mb | 2 | 0.0000 | 0.0058 | 0.0005 | 0.0158 | 0.0172 |
| breakpoints per 10 Mb | 3 | 0.0000 | 0.0000 | 0.0000 | 0.0030 | 0.0100 |
| absolute CN | 1 | 0.0000 | 0.0000 | 0.0038 | 0.0022 | 0.0013 |
| absolute CN | 2 | 0.0024 | 0.0032 | 0.0929 | 0.0922 | 0.0101 |
| absolute CN | 3 | 0.1778 | 0.1018 | 0.1453 | 0.1398 | 0.0456 |
| absolute CN | 4 | 0.0507 | 0.1234 | 0.0001 | 0.0313 | 0.1008 |
| absolute CN | 5 | 0.0044 | 0.0254 | 0.0000 | 0.0000 | 0.1117 |
| CN change point | 1 | 0.0356 | 0.1980 | 0.1388 | 0.2418 | 0.1516 |
| CN change point | 2 | 0.0003 | 0.0031 | 0.0009 | 0.0028 | 0.1065 |
| CN change point | 3 | 0.0000 | 0.0000 | 0.0000 | 0.0000 | 0.0052 |
| breakpoints per chr arm | 1 | 0.2123 | 0.0662 | 0.1239 | 0.0238 | 0.0004 |
| breakpoints per chr arm | 2 | 0.0000 | 0.0080 | 0.0013 | 0.0182 | 0.0140 |
| breakpoints per chr arm | 3 | 0.0000 | 0.0000 | 0.0000 | 0.0013 | 0.0040 |
| length of oscillating CN | 1 | 0.1994 | 0.0435 | 0.0942 | 0.0114 | 0.0478 |
| length of oscillating CN | 2 | 0.0006 | 0.0172 | 0.0083 | 0.0162 | 0.0278 |
| length of oscillating CN | 3 | 0.0000 | 0.0029 | 0.0003 | 0.0090 | 0.0099 |
| length of oscillating CN | 4 | 0.0000 | 0.0003 | 0.0000 | 0.0034 | 0.0001 |
| CN segment size | 1 | 0.0065 | 0.0447 | 0.0210 | 0.0338 | 0.0353 |
| CN segment size | 2 | 0.0077 | 0.0336 | 0.0175 | 0.0421 | 0.0529 |
| CN segment size | 3 | 0.0030 | 0.0259 | 0.0119 | 0.0483 | 0.0794 |
| CN segment size | 4 | 0.0011 | 0.0165 | 0.0147 | 0.0534 | 0.0615 |
| CN segment size | 5 | 0.0029 | 0.0420 | 0.0376 | 0.0495 | 0.0357 |
| CN segment size | 6 | 0.0261 | 0.0362 | 0.0362 | 0.0235 | 0.0047 |
| CN segment size | 7 | 0.0255 | 0.0192 | 0.0246 | 0.0116 | 0.0002 |
| CN segment size | 8 | 0.0618 | 0.0184 | 0.0384 | 0.0120 | 0.0002 |
| CN segment size | 9 | 0.0375 | 0.0048 | 0.0121 | 0.0000 | 0.0001 |
| CN segment size | 10 | 0.0659 | 0.0097 | 0.0252 | 0.0002 | 0.0002 |

**Supplementary Table 4. The sensitivity and specificity of chromothripsis prediction from copy-number (CN) signatures.** The probability of chromothripsis was obtained by receiver operator curve analysis using all CN signatures from each cohort as the input. Highlighted in green is the chromothripsis prediction probability of 0.6 which was used in the prediction model. (CN-SIG: copy-number signature from CoMMpass WGS; hCN-SIG: hematological cancer copy-number signature; eCN-SIG: copy-number signature from CoMMpass exome sequencing.)

| Prediction source | Probability | Sensitivity (%) | Specificity (%) |
|---|---|---|---|
| CN-SIG | **0.8** | 36 | 98 |
| | **0.7** | 48 | 97 |
| | **0.6** | 53 | 96 |
| | **0.5** | 58 | 93 |
| | **0.4** | 67 | 92 |
| | **0.3** | 75 | 89 |
| hCN-SIG | **0.8** | 34 | 98 |
| | **0.7** | 41 | 98 |
| | **0.6** | 48 | 98 |
| | **0.5** | 62 | 97 |
| | **0.4** | 69 | 96 |
| | **0.3** | 72 | 96 |
| eCN-SIG | **0.8** | 8 | 97 |
| | **0.7** | 16 | 97 |
| | **0.6** | 24 | 95 |
| | **0.5** | 34 | 91 |
| | **0.4** | 47 | 88 |
| | **0.3** | 59 | 82 |

**Supplementary Table 5. Summary clinical details of patients in the validation WGS dataset.**

| Clinical variable | Value |
|---|---|
| **Patients (n)** | 269 |
| **Gender (female, %)** | 40.9 |
| **Age (years, median, IQR)** | 59 (49-68) |
| **Cancer (n); Multiple Myeloma** | 34 |
| **Chronic Lymphocytic Leukemia** | 92 |
| **Chronic Myeloid Leukemia** | 29 |
| **Acute Myeloid Leukemia** | 10 |
| **B-Cell Lymphoma** | 104 |

IQR: interquartile range; n: number.

**Supplementary Table 6. A matrix of copy number (CN) category contributions defining 4 CN signatures (hCN-SIG) extracted from whole-genome sequencing of hematological patients in the validation dataset**

| CN feature type | Category code | hCN-SIG1 | hCN-SIG2 | hCN-SIG3 | hCN-SIG4 |
|---|---|---|---|---|---|
| breakpoints per 10 Mb | 1 | 0.0151 | 0.1999 | 0.1333 | 0.0281 |
| breakpoints per 10 Mb | 2 | 0.0000 | 0.0008 | 0.0080 | 0.0309 |
| breakpoints per 10 Mb | 3 | 0.0000 | 0.0000 | 0.0024 | 0.0056 |
| absolute CN | 1 | 0.0000 | 0.0047 | 0.0002 | 0.0030 |
| absolute CN | 2 | 0.0001 | 0.0695 | 0.0000 | 0.1261 |
| absolute CN | 3 | 0.2478 | 0.1008 | 0.0097 | 0.1640 |
| absolute CN | 4 | 0.0002 | 0.0431 | 0.0500 | 0.0003 |
| absolute CN | 5 | 0.0000 | 0.0030 | 0.2016 | 0.0008 |
| CN change point | 1 | 0.0001 | 0.1971 | 0.2160 | 0.2953 |
| CN change point | 2 | 0.0000 | 0.0000 | 0.0194 | 0.0001 |
| breakpoints per chr arm | 1 | 0.2465 | 0.0808 | 0.0334 | 0.0011 |
| breakpoints per chr arm | 2 | 0.0000 | 0.0000 | 0.0006 | 0.0011 |
| length of oscillating CN | 1 | 0.2402 | 0.0637 | 0.0698 | 0.0004 |
| length of oscillating CN | 2 | 0.0000 | 0.0024 | 0.0082 | 0.0246 |
| length of oscillating CN | 3 | 0.0000 | 0.0000 | 0.0002 | 0.0056 |
| length of oscillating CN | 4 | 0.0000 | 0.0000 | 0.0000 | 0.0006 |
| CN segment size | 1 | 0.0000 | 0.0172 | 0.0436 | 0.0568 |
| CN segment size | 2 | 0.0000 | 0.0189 | 0.0419 | 0.0768 |
| CN segment size | 3 | 0.0000 | 0.0261 | 0.0407 | 0.0773 |
| CN segment size | 4 | 0.0000 | 0.0288 | 0.0344 | 0.0659 |
| CN segment size | 5 | 0.0000 | 0.0464 | 0.0421 | 0.0298 |
| CN segment size | 6 | 0.0494 | 0.0788 | 0.0414 | 0.0055 |
| CN segment size | 7 | 0.0406 | 0.0125 | 0.0014 | 0.0000 |
| CN segment size | 8 | 0.0209 | 0.0052 | 0.0015 | 0.0001 |
| CN segment size | 9 | 0.0310 | 0.0000 | 0.0000 | 0.0000 |
| CN segment size | 10 | 0.1079 | 0.0001 | 0.0001 | 0.0000 |

**Supplementary Table 7. A matrix of copy number (CN) category contributions defining 5 CN signatures (eCN-SIG) extracted from whole-exome sequencing of newly diagnosed multiple myeloma patients in the CoMMpass dataset.**

| WGS CN signature | Validation CN signature | Validation cosine similarity | Exome CN signature | Exome cosine similarity |
|---|---|---|---|---|
| CN-SIG1 | hCN-SIG1 | 0.956 | eCN-SIG4 | 0.976 |
| CN-SIG2 | hCN-SIG2 | 0.920 | eCN-SIG2 | 0.962 |
| CN-SIG3 | hCN-SIG2 | 0.923 | eCN-SIG3 | 0.955 |
| CN-SIG4 | hCN-SIG4 | 0.943 | eCN-SIG1 | 0.915 |
| CN-SIG5 | hCN-SIG3 | 0.850 | eCN-SIG5 | 0.800 |

**Supplementary Table 8. Comparison between copy number signatures extracted from whole genome sequencing and those extracted from whole exome sequencing in newly diagnosed multiple myeloma.** CN-SIG: copy-number signature from CoMMpass WGS; hCN-SIG: hematological cancer copy-number signature; eCN-SIG: copy-number signature from CoMMpass exome sequencing.

| CN feature type | Category code | eCN-SIG1 | eCN-SIG2 | eCN-SIG3 | eCN-SIG4 | eCN-SIG5 |
|---|---|---|---|---|---|---|
| breakpoints per 10 Mb | 1 | 0.0249 | 0.1291 | 0.1417 | 0.0350 | 0.0055 |
| breakpoints per 10 Mb | 2 | 0.0192 | 0.0108 | 0.0004 | 0.0000 | 0.0229 |
| breakpoints per 10 Mb | 3 | 0.0000 | 0.0000 | 0.0000 | 0.0000 | 0.0000 |
| absolute CN | 1 | 0.0192 | 0.0018 | 0.0021 | 0.0000 | 0.0040 |
| absolute CN | 2 | 0.0807 | 0.0443 | 0.0374 | 0.0086 | 0.0545 |
| absolute CN | 3 | 0.1380 | 0.1121 | 0.1214 | 0.2311 | 0.0593 |
| absolute CN | 4 | 0.0562 | 0.0805 | 0.0648 | 0.0002 | 0.0098 |
| absolute CN | 5 | 0.0210 | 0.0252 | 0.0150 | 0.0000 | 0.1445 |
| CN change point | 1 | 0.2606 | 0.2232 | 0.1371 | 0.0049 | 0.0917 |
| CN change point | 2 | 0.0446 | 0.0099 | 0.0013 | 0.0000 | 0.0813 |
| CN change point | 3 | 0.0001 | 0.0004 | 0.0000 | 0.0000 | 0.0699 |
| breakpoints per chr arm | 1 | 0.0004 | 0.0287 | 0.1236 | 0.2418 | 0.0007 |
| breakpoints per chr arm | 2 | 0.0053 | 0.0170 | 0.0027 | 0.0000 | 0.0142 |
| breakpoints per chr arm | 3 | 0.0047 | 0.0003 | 0.0000 | 0.0000 | 0.0058 |
| length of oscillating CN | 1 | 0.0060 | 0.0260 | 0.1013 | 0.2339 | 0.1324 |
| length of oscillating CN | 2 | 0.0022 | 0.0181 | 0.0104 | 0.0000 | 0.0222 |
| length of oscillating CN | 3 | 0.0025 | 0.0081 | 0.0002 | 0.0000 | 0.0024 |
| length of oscillating CN | 4 | 0.0110 | 0.0014 | 0.0000 | 0.0000 | 0.0001 |
| CN segment size | 1 | 0.1117 | 0.0445 | 0.0328 | 0.0003 | 0.0799 |
| CN segment size | 2 | 0.0657 | 0.0338 | 0.0177 | 0.0001 | 0.0598 |
| CN segment size | 3 | 0.0432 | 0.0325 | 0.0081 | 0.0008 | 0.0669 |
| CN segment size | 4 | 0.0495 | 0.0427 | 0.0140 | 0.0000 | 0.0675 |
| CN segment size | 5 | 0.0267 | 0.0481 | 0.0228 | 0.0000 | 0.0022 |
| CN segment size | 6 | 0.0057 | 0.0430 | 0.0497 | 0.0254 | 0.0003 |
| CN segment size | 7 | 0.0002 | 0.0076 | 0.0159 | 0.0226 | 0.0002 |
| CN segment size | 8 | 0.0003 | 0.0099 | 0.0392 | 0.0683 | 0.0004 |
| CN segment size | 9 | 0.0002 | 0.0013 | 0.0212 | 0.0460 | 0.0015 |
| CN segment size | 10 | 0.0002 | 0.0000 | 0.0192 | 0.0809 | 0.0003 |
